# Functional stemness-related genes revealed by single-cell profiling of naïve and stimulated human CD34^+^ cells from CB and mPB

**DOI:** 10.1101/2022.02.23.481626

**Authors:** Guoyi Dong, Xiaojing Xu, Yue Li, Wenjie Ouyang, Weihua Zhao, Ying Gu, Jie Li, Tianbin Liu, Xinru Zeng, Huilin Zou, Shuguang Wang, Sixi Liu, Hai-Xi Sun, Chao Liu

## Abstract

Hematopoietic stem cells (HSCs) from different sources show varied repopulating capacity, and HSCs lose their stemness after long-time *ex vivo* culture. However, the underlying mechanisms of the stemness differences because of the cell sources and the culture stimulation are not fully understood. Here, we applied single-cell RNA-seq (scRNA-seq) to analyze the naïve and stimulated human CD34^+^ cells from cord blood (CB) and mobilized peripheral blood (mPB). We collected over 16,000 single-cell data to construct a comprehensive trajectory inference map and characterized the HSCs population on the hierarchy top, which is under quiescent state. Then we compared HSCs in CB to those in mPB and HSCs of naïve samples to those of cultured samples, and identified stemness-related genes (SRGs) associated with culture time (CT-SRGs) and cell source (CS-SRGs), respectively. Interestingly, CT-SRGs and CS-SRGs share genes enriched in the signaling pathways such as *mRNA catabolic process*, *Translational initiation*, *Ribonucleoprotein complex biogenesis* and *Cotranslational protein targeting to membrane*, suggesting dynamic protein translation and processing may be a common requirement for stemness maintenance. Meanwhile, CT-SRGs are enriched in pathways involved in glucocorticoid and corticosteroid response that affect HSCs homing and engraftment. In contrast, CS-SRGs specifically contain genes related purine and ATP metabolic process which is important to initiate hematopoiesis. Finally, we presented an application through a small-scale drug screening using Connectivity Map (CMap) against CT-SRGs and found a small molecule cucurbitacin I, targeting STAT3/JAK2, can efficiently expand HSCs *ex vivo* while maintaining its stemness. These results indicate SRGs revealed by scRNA-seq can provide helpful insights to understand the stemness differences under diverse circumstances, and CT-SRGs can be a valuable database to identify candidates enhancing functional HSCs expansion during *ex vivo* culture.

## INTRODUCTION

Hematopoietic stem cells (HSCs) are responsible for initiating hematopoiesis and maintaining the homeostasis of the hematopoietic system (Eaves, 2015), defined as cells on top of the hematopoietic hierarchy with totipotency (Notta et al., 2011). The molecular profile of human HSCs has been extensively investigated. Cell cycle senescence, glycolysis metabolism and self-renewal capacity are key biological signatures for human HSCs. Transcription factors, including *MEIZ1, TCF15* and *MLLT3*, and signal pathways, including Wnt, mTOR and HIF1, are involved in HSCs specification, differentiation and maintenance (Calvanese et al., 2019; Rodriguez-Fraticelli et al., 2020). Based on the above understanding, strategies aiming to modulate those factors or signaling pathways are explored to expand and maintain the functional HSCs *ex vivo* (Wilkinson et al., 2020). Meanwhile, unbiased screenings using small molecule libraries have been performed. Several candidates capable of efficiently expanding HSCs, including SR1 and UM171, have been identified and validated in animal models (Boitano et al., 2010; Fares et al., 2014). However, the strategies and small molecules mentioned above have not been successfully applied under clinical settings, suggesting other important underlying mechanisms required for HSC maintenance have not yet been identified.

In human, CD34^+^ cells isolated with magnetic beads containing the HSCs are clinically used for hematopoietic stem cell transplantation (HSCT) and hematopoietic stem cell gene therapy (HSC-GT). Increasing evidence shows that CD34^+^ cells are a heterogeneous cell population at different levels (Crisan and Dzierzak, 2016; Haas et al., 2018). Generally, to fully explore the HSC complexity, single-cell RNA sequencing (scRNA-seq) has been intensively used in this field with two approaches: one approach is to isolate the HSCs and other progenitor cells using fluorescence assisted cell sorting (FACS) with different antibodies against cell surface markers, and then perform the scRNA-seq in those “characterized” cell populations, respectively (Pellin et al., 2019); another approach is to perform scRNA-seq on unsorted CD34^+^ cells and then define the sub-cell populations based on bioinformatic analysis (Zheng et al., 2018). However, as the first approach still relies on the expression of surface markers, which may introduce bias in cell definition, it may not fully cover and reveal the *bona fide* cell populations when considerable cells are discarded during FACS process. Moreover, recent studies have found HSCs may also exist in other cell populations, suggesting the characterization based on cell surface markers are not sufficient. By contrast, the second approach, in theory, is able to capture all kinds of cells in an unbiased way, by which cell populations are clustered and defined by their gene expression profiles. However, when investigating cell sub-populations, particularly for HSCs, few known marker genes are reported and functional validation is lacking, making the reliability of these results questionable.

It is well-known that CD34^+^ cells derived from cord blood (CB) or mobilized peripheral blood (mPB) have different hematopoietic regeneration capacities (Mayani et al., 2020). For example in clinical practice, the minimal number of mPB CD34^+^ cells required for allogenic HSCT is usually above 2×10^6^/kg. By contrast for CB CD34^+^ cells, a quarter population (0.5×10^6^/kg) is sufficient for allogeneic cell transplantation (Remberger et al., 2015; Wagner et al., 2002). In addition, when generating humanized mice model, the required human CD34^+^ cells are 0.5×10^5^ of CB or 1.0×10^6^ of mPB cells per mouse(Wang et al., 1997), further indicating the higher stemness of CB than mPB CD34^+^ cells. On another aspect, the capacity of HSCs to rebuild the hematopoiesis is decreased along with the *ex vivo* culture time, as HSCs lose their stemness when they undergo activation and differentiation during the culture period (Glimm et al., 2000). Therefore, comparing CD34^+^ cells from CB and mPB sources, as well as from naive and cultured conditions would shed light on the signature genes and signaling pathways of HSCs.

Here we collected CB and mPB CD34^+^ cells from independent individuals, conducted the scRNA-seq immediately or after 48 hours *ex vivo* culture. We characterized HSCs population and identified two sets of stemness-related genes (SRGs) termed as CT-SRGs and CS-SRGs, associated with culture time and cell source, respectively. Using CT-SRGs to perform CMap searching, we found small molecule cucurbitacin I can efficiently expand HSCs *ex vivo* while maintaining its stemness. Our results demonstrate SRGs revealed by scRNA-seq can provide helpful insights for understanding the stemness maintenance of HSCs.

## RESULTS

### Identification of cell populations in human CD34^+^ cells

To characterize the human HSCs population and its functional signaling pathways, we collected CB and mPB CD34^+^ cells from fresh (termed naïve) or cultured (termed stimulated) conditions and captured their single-cell transcriptomes using a massively parallel single-cell library preparation technique, DNBelab C4 (Liu et al., 2019). With the comprehensive scRNA-seq dataset, we performed bioinformatic analysis to idendify HSC stemness related genes (SRGs) associated with culture time and cell source. Furthermore, we functionally validated the candidate small molecules predicted to regulate SRGs through *ex vivo* CD34^+^ cell culture and following FACS and scRNA-seq analysis. The overall experimental design is shown in Figure 1a (Fig. 1a).

**Figure 1.**
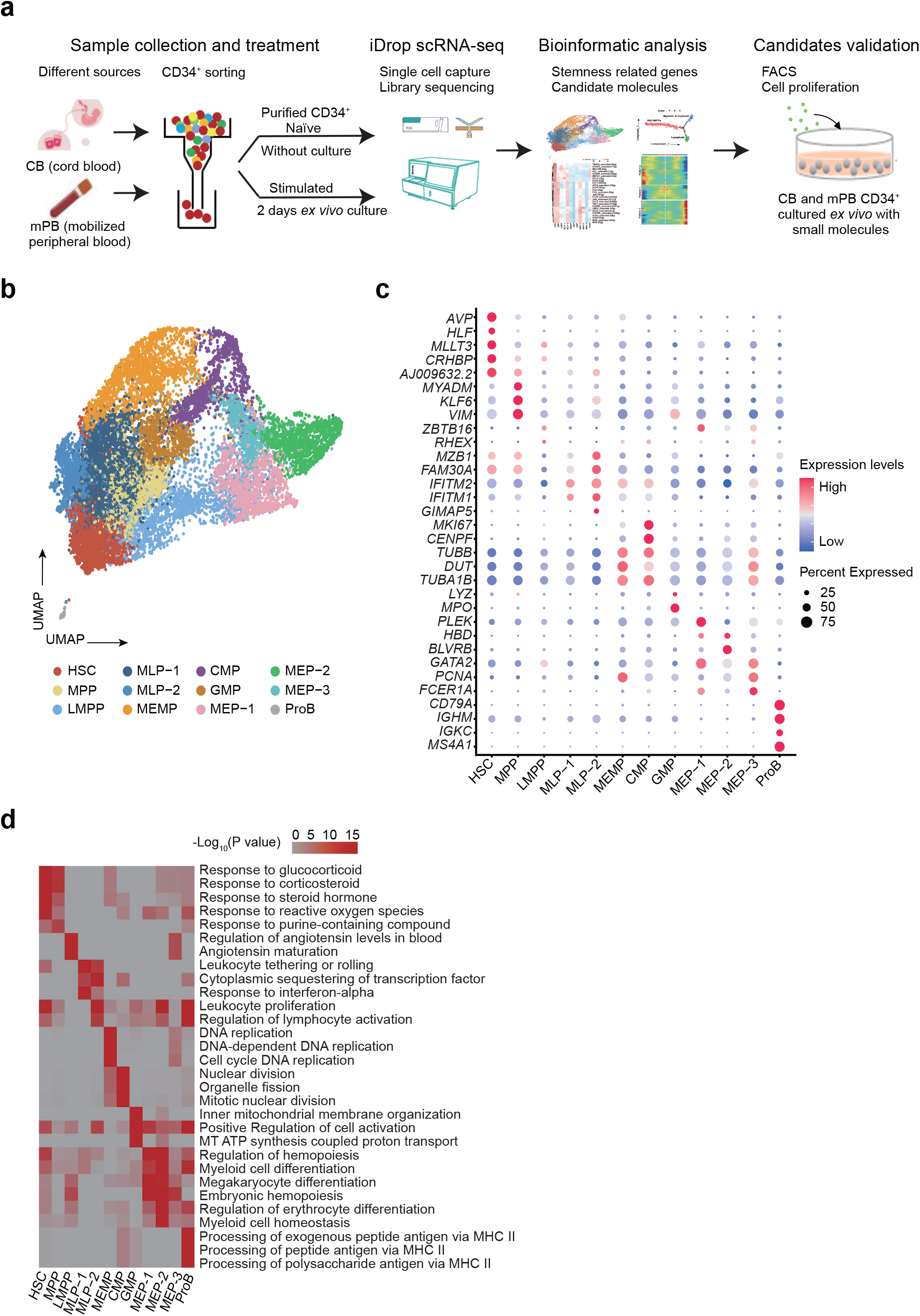
Single-cell transcriptome landscape of naïve and stimulated human CD34^+^ cells from cord blood (CB) and mobilized peripheral blood (mPB). a, Schematic diagram of the overall experimental design. b, CD34^+^ cells assigned to specific lineages by k-nearest neighbor (KNN) analysis are illustrated in the same UMAP space generated from the data. c, Dot plot displaying the expression of marker genes among of the identified 12 cell types. The node size positively correlates with the proportion of a given type of cells expressing a given marker gene. The color keys from blue to red indicate the range of relative gene expression in each cell type. d, Enrichment levels of the top enriched GO terms for marker genes of each cell type. The color keys from grey to red indicate the range of −log_10_-transformed *P* value.

We collected 2 biological replicates for each group, and after quality control we obtained 16,196 cells in total, with an average of ~3300 genes (~11,600 UMIs) per cell (Fig. S1a). Global correlation analysis revealed a strong correlation between biological replicates, and samples in the same culture conditions had higher transcriptome similarities than samples from the same source (Fig. S1b). To get an integrated single-cell transcriptome map of CD34^+^ cells, we performed graph-based clustering of the dataset, and found that almost all cells (99.8%, 16,171 of 16,196) were CD34 positive (Fig. S1c), which was consistent with our CD34^+^ cell sorting procedure (Fig. S1d). In summary, these results demonstrated that our transcriptome dataset had good quality for the subsequent analysis.

Next, we separated the cells into 12 populations and collected the top highly expressed marker genes to define cell types (Table. S1 and Table. S2). Based on these marker genes, we were able to assign the populations with distinct cell identities, including hematopoietic stem cells (HSCs), multipotent progenitors (MPPs), as well as myeloid, erythroid and lymphoid lineages (Fig. 1b). Many previously reported marker genes were also confirmed in our data (Fig. 1c, Fig. S2 and Table. S1), such as *AVP, MLLT3, HLF* and *CRHBP* of HSCs (Calvanese et al., 2019; Karamitros et al., 2018; Komorowska et al., 2017; Ranzoni et al., 2021; Xie et al., 2021; Zheng et al., 2018); *ZBTB16* of lymphoid-primed multipotent progenitors (LMPPs) (Karamitros et al., 2018); *TUBB, DUT* and *TUBA1B* of megakaryocyte-erythroid-mastcell progenitors (MEMPs) (Popescu et al., 2019); *MPO* and *LYZ* of granulocyte-monocyte progenitors (GMPs) (Karamitros et al., 2018; van Galen et al., 2019; Zheng et al., 2018); *HBD* and *GATA2* of megakaryocyte-erythroid progenitors (MEPs) (Karamitros et al., 2018; Zheng et al., 2018); *IGKC* and *MS4A1* of B cells progenitors (ProBs) (Popescu et al., 2019). To further validate the accuracy of cell type identification, we used a hypergeometric distribution test to evaluate the consistency between marker genes of cell clusters in our data and the top 500 up-regulated genes of cell types in eight published papers (Doulatov et al., 2010; Kohn et al., 2012; Laurenti et al., 2013; Laurenti et al., 2015; Lee et al., 2015; Milyavsky et al., 2010; Ng et al., 2009; Novershtern et al., 2011; Xie et al., 2021). By this analysis, we also observed a high consistency between these genes, providing further evidence supporting the cell type identification (Fig. S3).

Functional enrichment analysis of the marker genes further confirmed the characteristics of these cell types. For example, up-regulated genes in HSC/MPPs were related to cellular stress response as well as purine metabolic process, including *Response to glucocorticoid, Response to corticosteroid* and *Response to steroid hormone* that are reported as signature pathways enriched in HSCs in previous studies (Guo et al., 2017; Huang and Broxmeyer, 2019; Nakamura-Ishizu et al., 2020), whereas downstream progenitors were enriched for cell differentiation and cell activation related pathways in agreement with the cell development process and cell-cycle progression (Fig. 1d and Table. S3). Taken together, we obtained high-quality scRNA-seq data from 16,196 sorted CD34^+^ cells from naïve and stimulated CB and mPB, and identified 12 cell types including HSCs consistent with previous reports, providing a comprehensive reference map for investigating the underlying mechanism of stemness maintenance of HSCs.

### Differentiation trajectory of human HSCs

Previous studies found that HSCs first differentiated into MPPs, and then into LMPPs and other progenitor cells (Laurenti and Gottgens, 2018; Ng et al., 2009). Consistently, Principle Component Analysis (PCA) revealed that the cell types defined as adjacent developmental states in our map were clustered together, such as HSCs, MPPs and LMPPs which were all upstream progenitors (Fig. S4a). To further validate the accuracy of our reference map, we used Monocle (Trapnell et al., 2017) to conjecture the differentiation trajectory and checked whether our cell types exhibit similar pattern (Fig. 2a). Seven state cells were distributed along the trajectory and we found most HSCs/MPPs were located near the tips of the trajectory, while other cells were distributed amongst the six branches (Fig. 2b and Fig. S4b), in agreement with previous report. To determine the lineage affiliation of these branches, we checked the expression patterns of some marker genes. As we expected, the expression levels of *AVP, HLF* and *VIM*, all related to self-renewal potential and quiescence in HSC and MPP (Komorowska et al., 2017; Xie et al., 2021; Zheng et al., 2018), were decreased over the pseudo time. *ZBTB16* and *MZB1*, the marker genes of LMPP and MLP, were upregulated at the early stage but then dereceased later along the pseudo-time, in consistent with the position of LMPP and MLP on the trajectory. Myeloid and erythroid lineages such as MEMP, CMP, GMP and MEP were mainly situated at the end of the trajectory (State 5-7) with high expression of their lineage marker genes such as *DUT, CENPF, MPO* and *HBD* (Fig. 2c-d, Fig. S4b).

**Figure 2.**
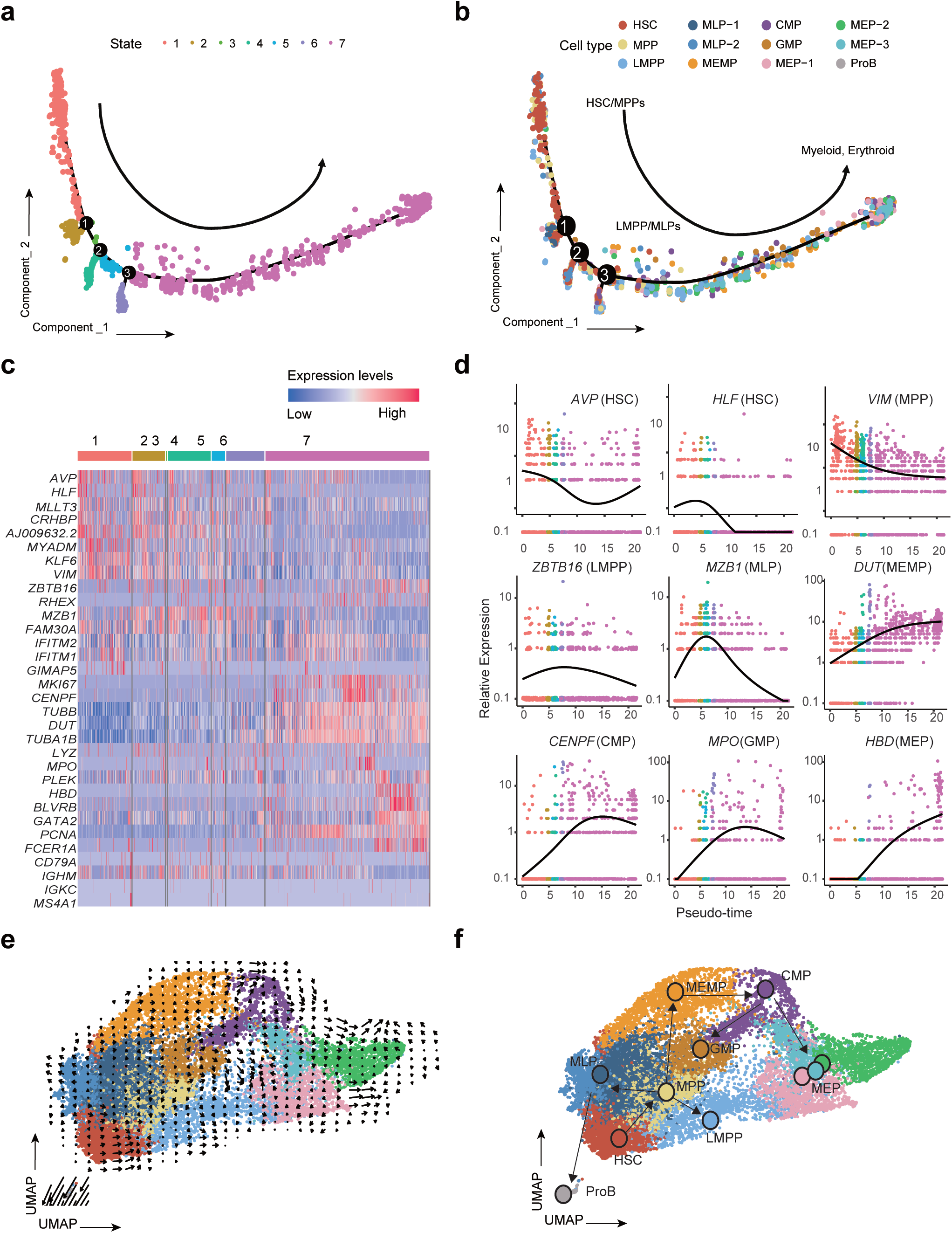
Developmental trajectory analysis of human HSCs. a, Developmental trajectory showing seven states for the 16,196 CD34^+^ cells. b, Cell type assignment along the developmental trajectory. HSCs/MPPs are mainly located in State1, LMPPs and MLPs lie at State2-4, myeloid and erythroid lineage cells gathered in State5-7. c, Heatmap showing the expression of marker genes (shown in Fig. 1c) in 7 developmental States. d, Expression of representative marker genes along developmental pseudo-time. e-f, amalgamated (e) and fitted (f) lineage trees showing the developmental routine from HSCs to multipotent progenitors and downstream myeloid, erythroid and lymphoid lineages. Arrows represent developmental directions, and each circle in (f) represents a cluster.

To further confirm the differentiation trajectory inferred, we used RNA velocity (La Manno et al., 2018) to project all cells into a two-dimensional UMAP space with arrows showing the direction and the speed of differentiation (Fig. 2e). When a developmental routine was finally fitted (Fig. 2f), we found that it was consistent with the current classical model of lineage determination in human hematopoietic hierarchy, thus further illustrating the accuracy of cell annotation and differentiation trajectory of our data.

### The characteristics and gene regulatory networks of human HSCs

To further characterize the human HSCs in our data, we did GO term analysis of the up-regulated genes of HSCs. We found pathways associated with cell cycle as well as mitosis, such as *regulation of spindle checkpoint, regulation of cell cycle spindle assembly checkpoint* and *mitotic cell cycle arrest*, were present as the most significant ones, which also includes other GO terms such as *response to glucocorticoid* and *response to corticosteroid* (Fig. 3a). The cell cycle activity of HSCs over the lifetime is dynamic, and it reflects the requirements of the organism at different developmental points (Gudmundsson et al., 2020; Lu et al., 2020). Thus, we calculated cell cycle phase scores based on canonical markers by Seurat (Stuart et al., 2019), and found that HSCs were significantly enriched in G0 and G1 phase (96.4%, *P* value = 5.62E-227, Fig. 3b and Table. S4) when compared with other cell types, indicating that most HSCs were in a resting state, in consistant with GO term analysis.

**Figure 3.**
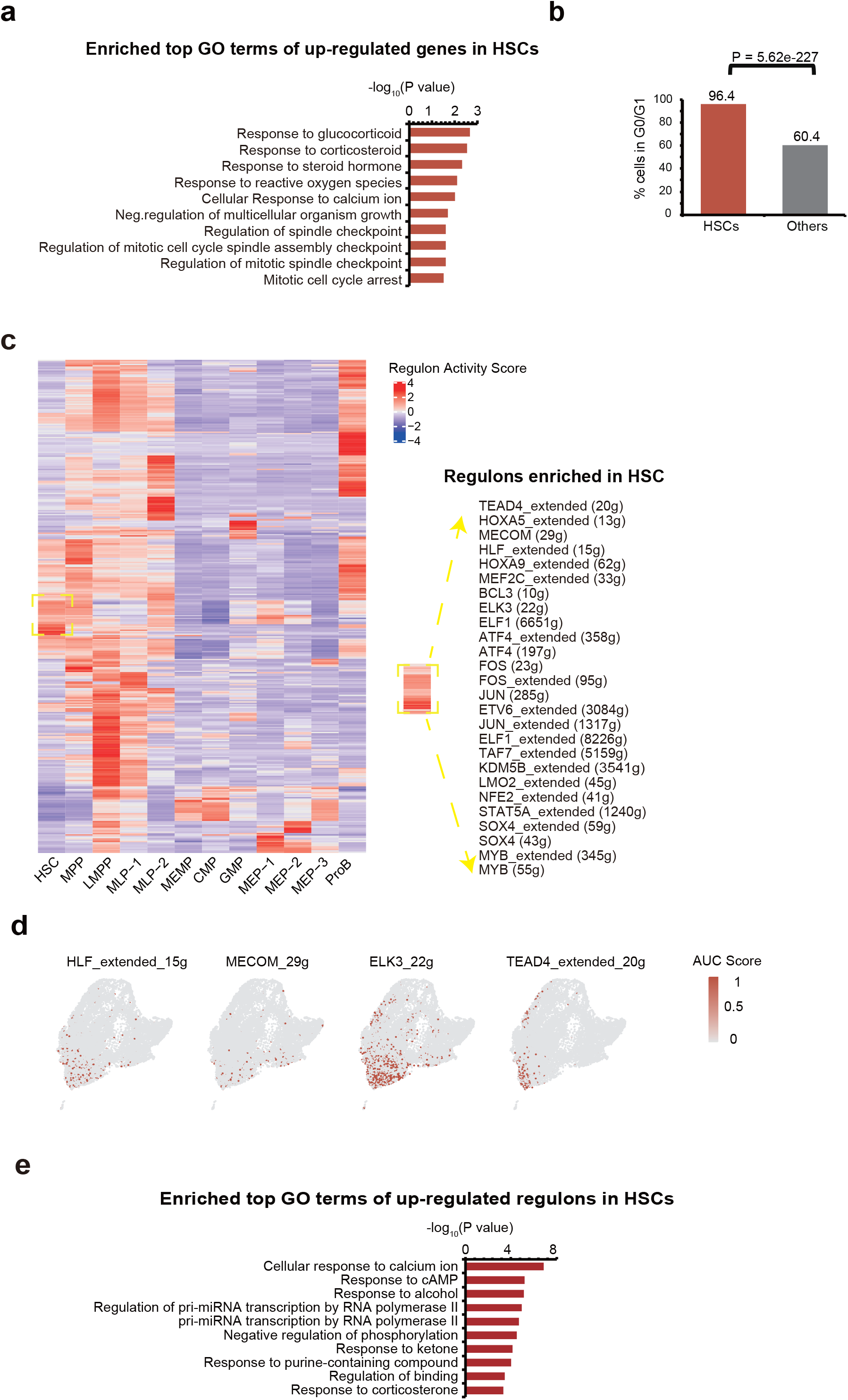
Cell cycle and gene regulatory networks of human HSCs. a, Enriched GO terms of up-regulated genes in HSCs compared with other cell types. b, The percentage of HSCs and other cells in G0/G1 stage. *P* value was calculated by Fisher’s Exact Test. c, Heatmap showing up-regulated regulons in each cell type compared to other cell types (left), up-regulated regulons in HSCs are listed in right. The value for each regulon is the regulon activity score. The color keys from blue to red indicate low to high SCENIC regulons activity levels in each cell type. d, UMAP plot showing the representative regulons of HSCs. Cells are colored by Area Under Curve (AUC) value of each regulon. The color keys from grey to red indicate the range of SCENIC TFs AUC score. e, Enriched GO terms of up-regulated regulons in HSCs.

Subsequently, we wondered whether some gene regulatory networks (regulons), a collection of genes regulated by common transcript factors (TFs), specifically existed in HSCs. To achieve this, we applied SCENIC (Aibar et al., 2017) to each cell of the 12 identified cell types. SCENIC recognized 371 activated regulons whose activities were dynamically changed among different cell types (Fig. 3c and Fig. S5). Interestingly, expression of TFs in several identified regulons, like *HLF, MECOM* and *ELK3*, were also enriched in HSCs and MPPs (Fig. 3c-d and Table S2). In addition, the functions of these TFs and their target gene sets were significantly enriched in *Response to corticosterone* and *Response to purine-containing compound* (Fig. 3e), which were consistent with those of HSC marker genes (Fig. 3a and Fig. 1d). Thus, using regulons analysis, we confirmed previous reported TFs, such as HLF and MECOM (Maicas et al., 2017), and their target genes, were also presented in HSCs of our data (Komorowska et al., 2017; Zheng et al., 2018). Besides, we also found many novel regulons worthy of further investigations, including *FOS, MEF2C*, *TEAD4, ELK3* and *HOXA9* (Fig. 3c-d).

### Identification of stemness-related genes (SRGs) probably controlling stemness of human HSCs

HSCs accounted for 9.9% of all cells profiled in our data, and we found that their composition was significantly decreased under stimulated condition in both CB and mPB samples (Fig. 4a), which was in agreement with the reduced stemness of these samples. Interestingly, we noticed that cell sources and 2 days *ex vivo* culture didn’t affect the cell cycle phase scores (Fig. 4b), suggesting quiescent maintenance is a robust characteristic for all HSCs, even after short time *ex vivo* culture. To reveal SRGs responsible for stemness maintenance of HSCs, we investigated the transcriptome changes between naïve and stimulated samples. By differential expression analysis, we obtained 247 CT-SRGs (Culture Time-related SRGs), from the intersection of 716 up-regulated genes in CB CD34^+^ naïve and 1,164 up-regulated genes in mPB CD34^+^ naïve (*P* value < 0.05 and ln-transformed fold change > 0.25) (Fig. 4c-e and Table. S5-7). In addition, as previous studies found that CD34^+^ cells derived from CB exhibited a higher level of stemness than mPB (Eliane et al., 2011; Martínez-Jaramillo et al., 2004; Wang et al., 2003), we also obtained 560 CS-SRGs (Cell Source-related SRGs) by comparing naïve CB and naïve mPB (*P* value < 0.05 and ln-transformed fold change > 0.25) (Fig. 4f and Table. S8). Gene set variation analysis (GSVA) using HSC data sets from two published papers (Milyavsky et al., 2010; Novershtern et al., 2011) revealed that the overall expression levels of both CT-SRGs and CS-SRGs were significantly higher than HSC lineage distinct genes (HSC-LDGs) (Fig. 4g-h), further demonstrating that both CT-SRGs and CS-SRGs could better reflect the stemness of HSCs.

**Figure 4.**
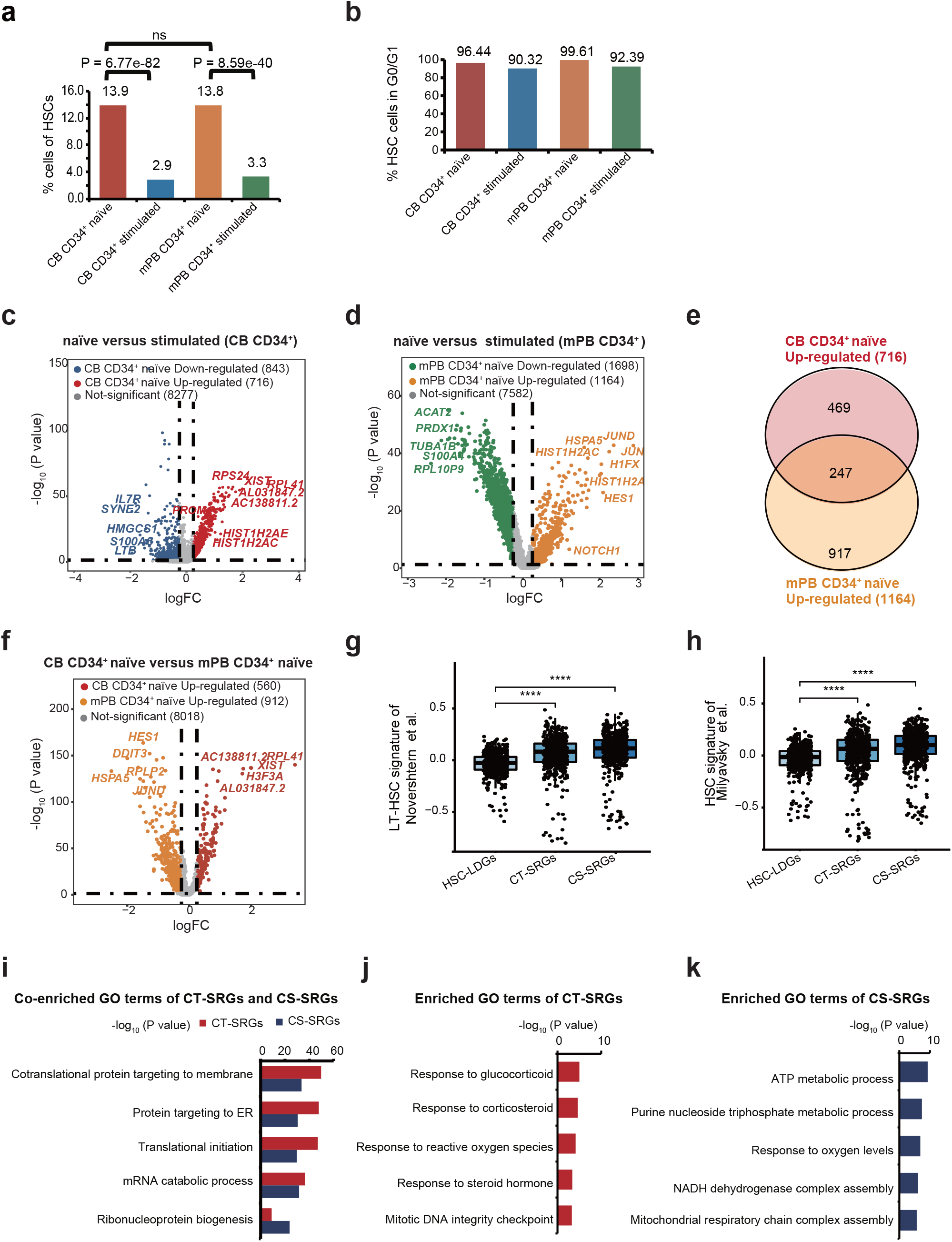
Changes in the transcriptional profiles after stimulation of HSCs during human hematopoiesis. a, Proportion of HSCs in naïve and stimulated CD34^+^ cells from CB and mPB. *P* value was calculated by Fisher’s Exact Test. ns means not significant. b, The percentage of HSC cells in G0/G1 stage per sample. *P* value was calculated by Fisher’s Exact Test, There was no difference between the two samples. c, Volcano plots showing gene expression changes of HSCs between naïve CB and stimulated CB. d, Volcano plots showing gene expression changes of HSCs between naïve mPB and stimulated mPB. e, Venn diagram showing up-regulated genes in naïve compared with stimulated samples of HSCs between CB and mPB. f, Volcano plots showing gene expression changes of HSCs between naïve CB and naïve mPB. g-h, Violin plots showing the scores of HSC marker genes (HSC-LDGs), CT-SRGs and CS-SRGs of HSCs identified in Novershtern *et al*. (g) and Milyavsky *et al*. (h). The horizontal axis represents different gene sets, and the vertical axis represents GSVA enrichment scores of signature genes of HSCs in the paper. *P* value was calculated by two-sided Wilcoxon rank-sum test. i, Common enriched GO terms of CT-SRGs and CR-SRGs. GO terms enriched in CT-SRGs are shown in red, whereas GO terms enriched in CS-SRGs are shown in blue, the colored bars indicate the range of −log_10_-transformed *P* value. j, Specifically enriched GO terms of CT-SRGs compared to CS-SRGs. k, Specifically enriched GO terms of CS-SRGs compared to CT-SRGs.

Then we asked the commonalities and differences between CT-SRGs and CS-SRGs. Interestingly, a large fraction of the intersection genes in CT-SRGs and CS-SRGs are enriched in GO terms related to *Protein targeting to membrane* (P value (CT-SRGs) = 1.94E-50, P value (CR-SRGs) = 2.25E-34) and *Protein targeting to ER* (P value (CT-SRGs) = 1.48E-48, P value (CR-SRGs) = 3.26E-31) (Fig. 4i), which are signatures for protein synthesis process and translation of ribosomal coding genes. Next, we It is worth noting that nine genes specifically present in CT-SRGs, including *AREG, CFLAR, DDIT4, DUSP1, FLT1, FOS, FOSB, FOXO3* and *ZFP36*, were significantly enriched in GO terms of *Response to glucocorticoid* (P value = 8.50E-06) and *Response to corticosteroid* (P value = 1.96E-05) (Fig. 4j). By contrast, we found *“Purine nucleoside triphosphate metabolic process* (P value = 4.90E-12) and *ATP metabolic process* (P value = 4.48E-10) were specifically enriched in CS-SRGs (Fig. 4k). Taken together, these results demonstrated that both CT-SRGs and CS-SRGs exhibited a better consistency with HSCs characteristics than HSCs marker genes, indicating their potential applications in stemness maintenance of human HSCs.

### The differentiation trajectory of human CD34^+^ cells reconstructed using CT-SRGs

During HSCT and HSC-GT, the ability of CD34^+^ cells to rebuild the hematopoiesis is negatively correlated with the *ex vivo* culture time, therefore to understand the transcriptomic changes during this process, we focused on CT-SRGs in further analysis. First, we asked whether we can reconstruct the differentiation trajectory of human CD34^+^ cells using CT-SRGs (Fig. 5a). Consistent with the trajectory conjectured based on marker genes (Fig. 2), HSCs/MPPs were still located at the top of the trajectory. Additionally, we were amazed to find that lymphoid lineage (State2) and myeloid lineage (State3) were clearly separated in the developmental trajectory after HSCs/MPPs (State1) (Fig. 5b and Fig. S6a), indicating that CT-SRGs had different expression patterns when HSCs differentiate into these two lineages. We further checked the expression changes of marker genes along pseudo-time. *AVP*, *HLF, VIM* and *KLF6* of HSCs/MPPs were highly expressed in State1, then decreased with the pseudo-time progression. Oppositely, *MKI67* and *HBD* gradually increased with pseudo-time and reached the peak in State3, signifying myeloid and erythroid lineages may enrich in State3. State2 might be lymphoid lineage with high expression of *IGKC* and *MS4A1* (Fig. S6b). Next, we identified differentially expressed genes between the branches to further corroborate the previous results, and we got three gene clusters with different expression patterns across three States. GO Term analysis of the three gene clusters indicated related pathways were enriched in corresponding states, such as *lymphoid differentiation* in Lymphoid, *cotranslational protein targeting to membrane* in HSCs/MPPs and *neutrophil activation* in Myeloid&Erythroid, further confirming the reliability of these results (Fig. 5c). A total of 15 CT-SRGs were up-regulated in State2 (lymphoid progenitors), whereas no CT-SRG was up-regulated in State3 (Fig. 5d), indicating that lymphoid progenitors may be more close to HSCs when compared to myeloid or erythroid cells. In agreement, the 15 CT-SRGs upregulated in lymphoid were also highly expressed in MPPs and HSCs (Fig. 5e). In conclusion, CT-SRGs may be better reference genes for development trajectory construction, revealing a closer relationship between HSCs and lymphoid lineage.

**Figure 5.**
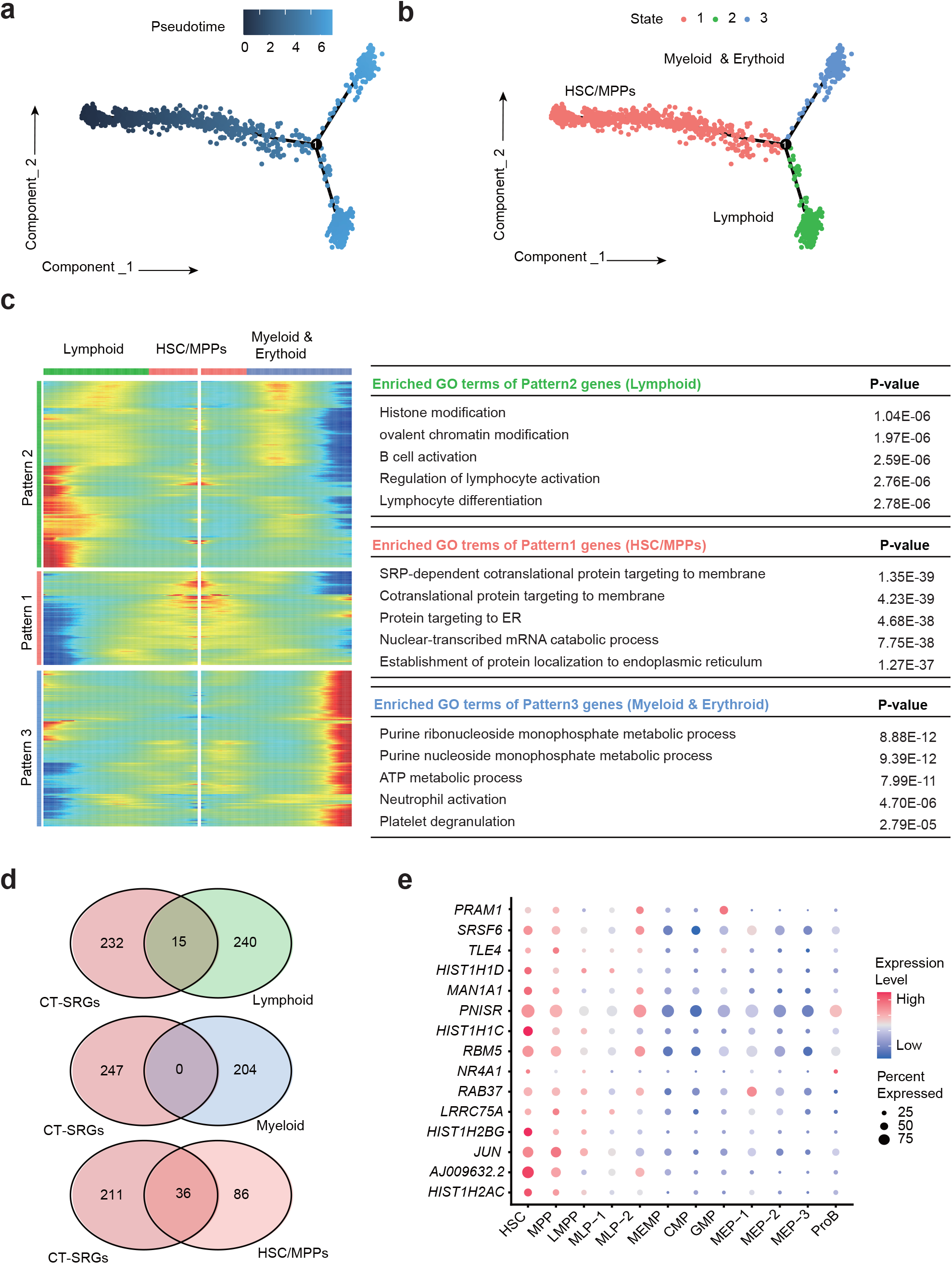
Differentiation and development trajectory based on CT-SRGs. a, The trajectory tree based on pseudo time staining. b, The State branch recognized by Monocle2. c, Heatmap of the key genes involved in branch determination and their functions. Heatmap showing the three dynamic gene expression patterns of Lymphoid, HSCs/MPPs and Myeloid & Erythoid (left). The specifically enriched GO terms are shown on the right. d, Venn diagram showing shared genes between CT-SRGs and three clusters (shown in c). e, Dot plot displaying the expression of 15 shared genes in CT-SRGs and cluster1 (Lymphoid). The node size represents the cell proportion that expresses the given gene. The color keys from blue to red indicate the range of relative gene expression.

### Small molecules modulating CT-SRG expression promote cell proliferation and stemness maintenance of human HSCs *ex vivo*

Connectivity Map (CMap) is an online tool kit based on a perturbation-driven gene expression dataset (Lamb et al., 2006). To functionally validate the effectiveness of CT-SRGs in maintaining stemness of human HSCs, we used CMap to search for candidate small molecules that could affect the global expression levels of CT-SRGs and then performed the experimental validation process (Fig. 6a). We identified 145 candidates in total. Among them, small molecules function as protein synthesis inhibitor and glucocorticoid/corticosteroid receptor agonist were predicted to positively regulate the expression levels of CT-SRGs, consistent with the above results showing that the functions of CT-SRGs were enriched in protein synthesis process and glucocorticoid/corticosteroid responses. In addtion, small molecules that target ATPase, mTOR and MAP kinase pathways, were also characterized as candidates in our screening (Table S9). Importantly, fenretinide, a retinoid receptor agonist and identified in our CMap screening, has been reported to enhance human HSCs self-renewal by modulating sphingolipid metabolism *ex vivo* (Xie et al., 2019), demonstrating small molecules predicted to modulate CT-SRGs can be candidates capable of regulating human HSCs stemness.

**Figure 6.**
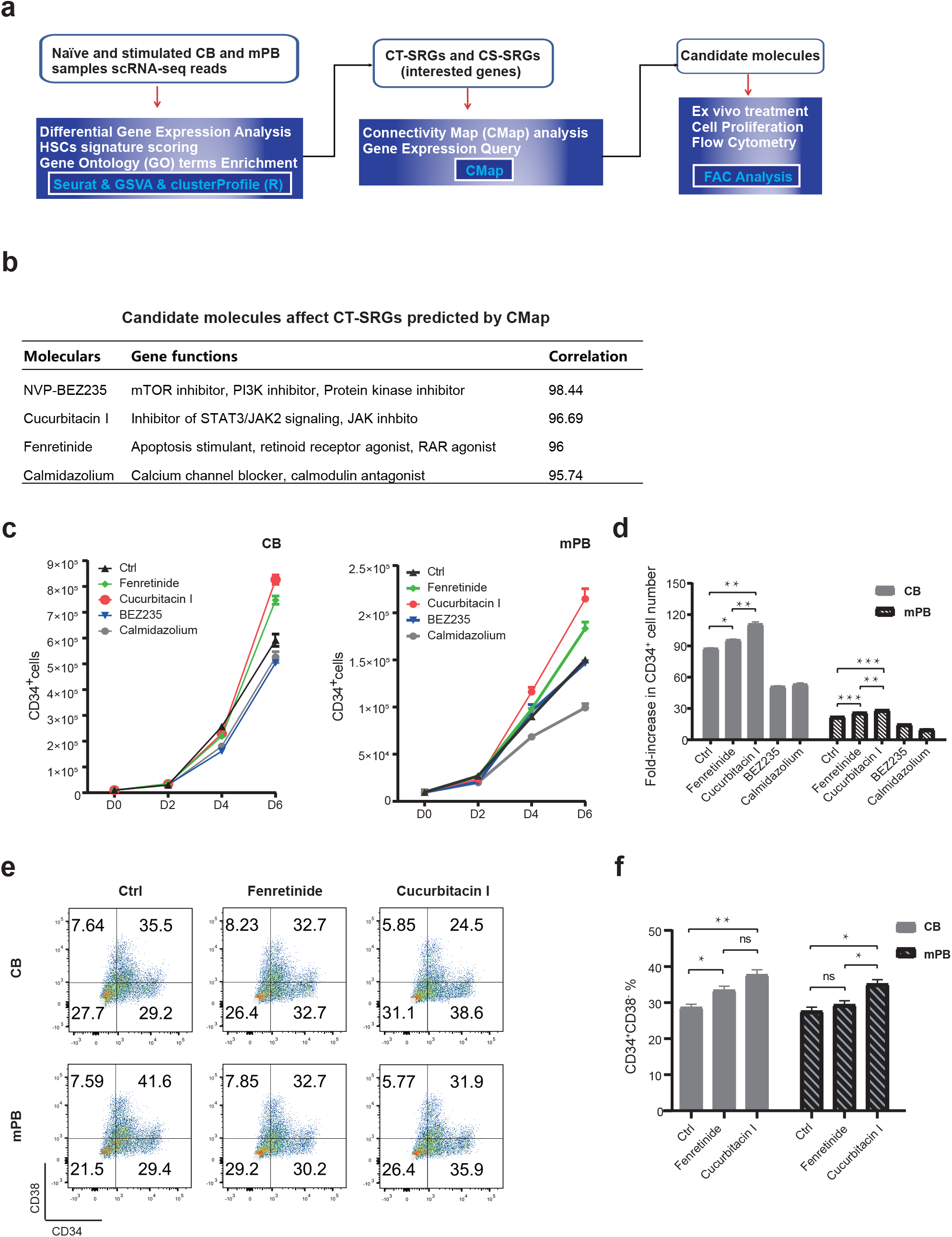
Cucurbitacin I could enhance HSCs proliferation and stemness. a, The small molecules screening pipeline. b, Candidate molecules affect CT-SRGs predicted by CMap. c, Proliferation kinetics of CD34^+^ cells from CB and mPB in candidate small molecules culture. d, Fold-increase in CD34^+^ cell number on 6 days culture ex vivo as compared to input numbers. e, Flow cytometry plots show CD34^+^CD38^-^ populations on 6 days culture ex vivo from CB and mPB. f, Proportion of CD34^+^CD38^-^ cells after different small molecule treatments detected by Flow Cytometry. P value was calculated by *t*-test, *p < 0.05; **p < 0.01; ***p < 0.001; ns, not significant. Error bars indicate standard deviations of triplicate cultures. For fenretinide and cucurbitacin I treatment, 100 nM dose was used. f,

To identify other regulators of HSCs stemness, we selected NVP-BEZ235, Cucurbitacin I and Calmidazolium as these molelues were related to the signal of mTOR inhibitor, PI3K inhibitor, Protein kinase inhibitor, Inhibitor of STAT3/JAK2 signaling, JAK inhibitor, Calcium channel blocker, calmodulin antagonist (Fig. 6b). We treated CB and mPB CD34^+^ cells *ex vivo* with those small molecule candidates, cultured them for extended time period and then measured the cell proliferation and HSCs percentage. Fenretinide was used as a positive control. When compared to fenretinide treatment and untreated control, only Cucurbitacin I, but not other tested molecules, increased the CD34^+^ cell numbers in CB and mPB after 6 days’ *ex vivo* culture (Fig. 6c-d). More important, when examining a more restricted surface markers of HSCs using FACS, the cucurbitacin I treated cells also exhibited the highest proportion of CD34^+^CD38^-^ cells, indicating that cucurbitacin I not only enhances the expansion but also promote human HSCs maintenance of both CB and mPB sources (Fig. 6e-f). Taken together, we provided a preliminary application of CT-SRGs to identify HSCs modulators and validated that small molecule cucurbitacin I could enhance cell proliferation and stemness maintenance of human HSCs.

## DISCUSSION

Many previous studies have demonstrated that using scRNA-seq transcriptomic data, a marker-free approach, can capture detailed molecular characterization of single cells and unbiased define cell clusters during hematopoiesis (Giladi et al., 2018; Zheng et al., 2018). Using this method, many novel cell types have been found and functional identification (Villani et al., 2017). In our present study, we used scRNA-seq data from CD34^+^ cells to accurately identify all classical human hematopoietic stem and progenitor cells (HSPCs) cell types reported in previous researches (Laurenti and Göttgens, 2018b), including HSCs, without prior specific cell surface markers sorting. It is worth mentioning that our data shows better data quality, being embodied in a strong correlation between biological replicates and a more credible cell type identification. Based on these molecular data, we found that most HSCs stay in G0 and G1 phase, indicating that cells were in a quiescent state, those critical transcription factors for the maintenance and proliferation of HSCs, such as *AVP*, *MLLT3*, *HLF*, *MECOM*,*CD52*, are highly expressed.

HSCs from CB and mPB process distinct ability of reconstruction of hematopoiesis and loss stemness *in vitro* culture. Uncovering the molecular distinction is helpful to find out the underlying mechanisms of HSCs maintenance and proliferation. We roundly compared HSCs in CB to those in mPB and HSCs of naïve samples to those of cultured samples, and identified SRGs associated with culture time (CT-SRGs) and cell source (CS-SRGs), responsible for stemness maintenance and hematopoiesis initiation, respectively. We found that naive CB CD34^+^ may be able to both rapidly colonize and maintain stemness, whereas mPB CD34^+^ may tends to be naive CB CD34 colonization state with activation on the one hand, and loses glucorticoid pathway compared to naive CB CD34^+^ on the other hand. Meanwhile, we verified the reliability of the method to screen stemness-related genes.

One obstacle limiting the wide application of HSCs in clinics is the lack of effective methods to culture and expand HSCs *ex vivo* while maintaining its stemness. Thus we focused on investigating CT-SRGs and identified new molecules using CMap analysis. Interestingly, we found that small molecule cucurbitacin-I could boost cell proliferation and stemness maintenance of human HSCs. Cucurbitacin I is a STAT3/JAK2 inhibitor, however, its function in human HSCs expansion and stemness maintenance has not been clarified. In this study, we suggest CucurbitacinI may be a novel therapeutic molecule to expanding HSCs for HSCT and HSC-GT in future clinical applications.

## METHODS

### Enrichment of CD34^+^ cells from human CB and mPB samples

Human CB and mPB samples were obtained with informed consent from XXX. Mononuclear cells (MNC) were obtained by centrifugation on Lymphoprep medium, and after ammonium chloride lysis MNC was enriched for CD34^+^ cells by positive selection with the CD34 Microbead kit and LS column purification with MACS magnet technology (Miltenyi). The sorted CD34^+^ cells were subject to downstream experiments.

### Cell cultivation and scRNA-seq

Fresh CD34^+^ cells were immediately cultured in vitro or single-cell RNA-seq. For cell cultivation, CD34^+^ cells were resuspended in SCGM medium (Cellgenix) with following recombinant hematopoietic cytokines: Recombinant Human stem cell factor (rhSCF) 100ng/ml, Recombinant Human Thrombopoietin (rhTPO)100ng/ml, Recombinant Human Fms-related Tyrosine Kinase 3 Ligand (rhFlt3-L) 100ng/ml,, and cultured in 24-well tissue culture plates at 37°C in an atmosphere of 5% CO_2_ in the air for 48 hours and collected to scRNA-seq. For scRNA-seq were performed by the DNBelab C4 platform (Liu et al., 2019). In brief, single-cell suspensions were used for droplet generation, emulsion breakage, beads collection, reverse transcription, and cDNA amplification to generate barcoded libraries. Indexed libraries were constructed according to the manufacturer’s protocol. The sequencing libraries were quantified by QubitTM ssDNA Assay Kit (Thermo Fisher Scientific, #<u>Q10212</u>). DNA nanoballs (DNBs) were loaded into the patterned nano arrays and sequenced on the ultra-high-throughput DIPSEQ T1 sequencer using the following read length: 30 bp for read 1, inclusive of 10 bp cell barcode 1, 10 bp cell barcode 2, and 10 bp unique molecular identifier (UMI), 100 bp of transcript sequence for read 2, and 10 bp for sample index.

### Quality control of scRNA-Seq data

The iDrop Software Suite (v.1.0.0) was applied to perform sample demultiplexing, barcode processing, and single-cell 3’ unique molecular identifier (UMI) counting with default parameters. Processed reads were then aligned to the complete UCSC hg38 human genome by splicing-aware aligner STAR (Dobin et al., 2013) with default parameters. Gene-cell metrics were generated for advanced analysis of valid cells that were automatically recognized according to the UMI number distribution of each cell.

The R (v.3.6.3) package Seurat (v.3.2.1) (Butler et al., 2018; Stuart et al., 2019) was used to perform the following steps: 1) Quality control of three indicators: the number of genes expressed per cell, the number of UMI (Unique Molecular Identifiers) and the proportional distribution of mitochondrial RNA to screen high-quality cells for subsequent analysis(Griffiths et al., 2018; Ilicic et al., 2016). Due to the differences among samples, for example, cryopreserved (naïve) and stimulated, the number of genes expressed varied greatly, thus we selected Tukey’s test method (Kannan et al., 2015) to remove cells with abnormal gene numbers. Cells that expressed genes lower than Q1-IQR or higher than Q3+IQR were removed. Meanwhile, cells with a mitochondrial mRNA ratio greater than 10% were also removed; 2) Doublets removal. We used an R package DoubleFinder (v.2.0.3) (McGinnis et al., 2019) to remove doublets; 3) Batch effect removal. We created an integrated data assay of all samples by identifying anchors using FindIntegrationAnchors function; 4) Data normalization was performed using NormalizeData function with scaling factor 10,000 and then log-transformed the data. 5) Detection of 4,000 highly variable genes (HVGs) by FindVariableFeatures function with “vst” method; 6) Scaling of the features by ScaleData function to get a unit variance and zero mean of all samples.

### Dimensionality reduction and cell cluster

We perform PCA on the previously determined variable features after scaled, by RunPCA function with top 40 significant PCs that represent a robust compression of the dataset. Next, we applied a graph-based clustering approach to construct a shared nearest neighbor graph for a given dataset by FindNeighbors function between every cell and optimize the modularity function to determine clusters by FindClusters function with resolution equal to 0.6. Finally, we used UMAP to learn the underlying manifold of the data and place similar cells together in low-dimensional space.

### Cell type annotation

We executed biologically-related pairwise differential gene expression analysis between duos of clusters by FindAllMarkers function with min.pct equal to 0.2 and logfc.threshold set to 0.25, to identify DEGs and to examine the quantitative changes in the expression levels between the clusters.

Because the marker genes of HSCs and their downstream progeny cells are still uncertain, we additionally collected nine bulk RNA-seq datasets to improve the cell definition. Detailed procedures are as follows: 1) Using GEO2R, a NCBI online tool, to calculate gene expression levels of reference datasets, respectively; 2) Selected top 500 significantly up-regulated genes as biomarkers of each cell type; 3) Performed Hypergeometric distribution test between differentially expressed genes of each dataset and current data, and assigned the cell type based on the significance of *P* value.

### Differential gene expression analysis

We executed biologically-related pairwise differential gene expression analysis between duos of samples to identify DEGs and to examine the quantitative changes in the expression levels between the samples in HSCs. We calculated the DEGs by applying the FindMarkers function (Wilcoxon rank-sum with adjusted *P* values for multiple testing with the Benjamini-Hochberg correction). We filtered out the obtained DEGs by setting min.pct to 0.2, so that a gene is expressed in at least 20% of the cells in one of the two tested groups. A gene was considered significant with adjusted P < 0.05 and logFC > 0.25.

### RNA velocity analysis

By distinguishing between unspliced and spliced mRNAs, RNA velocity, the time derivative of the gene expression state, can be directly estimated. Thus, we utilized velocyto (La Manno et al., 2018) to compute the rate of transcriptional alteration of each cell.

### Developmental trajectory inference analysis

The Monocle2 (v.2.14.0) (Qiu et al., 2017; Trapnell et al., 2017) algorithm with the core SRGs was applied to order all cells in pseudo time. By creating an object with parameter “expressionFamily = negbinomial.size”, regressed out the batch effect using the “reduceDimension” function with residualModelFormulaStr setting to exclude technology influence and with default reduction_method to achieve dimensionality reduction. Cell differentiation trajectory was determined successfully built based on the above steps.

Next, the BEAM function was used to detect genes that separate cells into the considered cell branches. We used the plot_multiple_branches_heatmap function to separate the branch-related gene set with a q-value less than or equal to 10e^-4^ and the num_clusters = 3.

### Gene Ontology (GO) terms enrichment analysis

GO enrichment analysis was performed on the given gene set. The enrichGO function of the “clusterProfiler (Wu et al., 2021)” R package was used to do enrichment analysis. Terms with the q value < 0.05 corrected by FDR were considered statistically significant.

### Prediction of transcription factor regulons

To predict transcription factor regulons, we utilized SCENIC (v.1.1.3) (Aibar et al., 2017; Van de Sande et al., 2020) with default parameters and used cisTarget database (https://resources.aertslab.org/cistarget/). Generated AUC scores of cells were applied for downstream analysis.

### Connectivity Map (CMap) analysis

We used Gene Expression (L1000) in Query (https://clue.io/query, a tool of CMap (Lamb et al., 2006)) with default parameters to compare the similarity between the two SRG signature lists and the expression profile in the reference database.

### HSCs signature scoring

We collected the top 250 high expressed genes as HSCs marker genes of each two published papers (Milyavsky et al., 2010; Novershtern et al., 2011), utilizing GSVA(Hanzelmann et al., 2013) to score each HSCs cell, respectively. Meanwhile, we scored each HSCs cell by HSCs marker genes in current data.

### FAC analysis

Cells cultured with selected small mollecus were collected at day 6, washed in DPBS, and then incubated with antibody CD34(Biolegend), CD38 (BD) at 4°C for 30min, washed and resupended in DPBS for FAC analysis.

## DATA ACCESS

The scRNA-seq data generated in this study have been deposited in the CNSA (https://db.cngb.org/cnsa/) of CNGBdb with accession code CNP0000978.

## ETHICS APPROVAL

Permission for this study was obtained from the Bioethics and Biological Safety Review Committee of BGI-Shenzhen (BGI-IRB 20155). We confirm that all experiments were performed in accordance with relevant guidelines and regulations.

## ACKNOWLEDGMENTS

We thanks the support provided by China National GeneBank, and this research is supported by grants from National Natural Science Foundation of China (NSFC) (31970816).

## AUTHOR CONTRIBUTIONS

G.D., S.L., H.S., and C.L. conceived and designed the study. W.Z., Y.L., S.W., and S.L. collectied the CB and mPB samples. G.D., W.O., T.L., X.Z. and H.Z. performed the experiments. X.X., J.L. and H.S. conducted the bioinformatics analysis. Y.G. helped project design and discussions. G.D. and X.X.wrote the manuscript. H.S. and C.L. reviewed the manuscript. All authors read and approved the final manuscript.

---

**Figure S1. Quality control and cell type identification of human hematopoietic stem cells (HSCs).**

a, Violin plots showing the distribution of gene number (left), UMI number (middle) and percentage of mitochondrial genes (right) of 16,196 cells. b, Hierarchical clustering of all sample correlations from CB and mPB under naïve and stimulated conditions. Pearson correlation coefficient (PCC) is represented by the color-coded scales on the right. c, UMAP visualization of *CD34* expression. Color scale indicates gene expression. d, FACS analysis of CD34 marker of naïve CB/mPB. The horizontal axis represents the intensity of the fluorescence signal of the CD34 marker, and the vertical axis represents the number of CD34^+^ cells.

**Figure S2. Expression of marker genes of identified cell types.**

a. Expression of HSC markers (*AVP, MLLT3, HLF, CRHBP*, and *AJ009632.2*). b. Expression of MPP markers (*VIM, KLF6* and *MYADM*). c. Expression of LMPP markers (*RHEX* and *ZBTB16*). d. Expression of MLP markers (*MZB1, IFITM1* and *GIMAP5*). e. Expression of ProB markers (*MS4A1* and *IGKC*). f. Expression of MEMP markers (*TUBA1B, DUT* and *TUBB*). g. Expression of CMP markers (*MKI67* and *CENPE*). h. Expression of GMP markers (*MPO* and *LYZ*). i. Expression of MEP markers (*HBD, BLVRB* and *FCER1A*). Color scale indicates gene expression.

**Figure S3. Proof of cell type allocation.**

a, Heatmap of enrichment score between cell types identified in current data and the data of (Laurenti et al., 2013). b, Heatmap of enrichment score between cell types identified in current data and the data of (Doulatov et al., 2010). c, Heatmap of enrichment score between cell types identified in current data and the data of (Novershtern et al., 2011). d, Heatmap of enrichment score between cell types identified in current data and the data of (Kohn et al., 2012). e, Heatmap of enrichment score between cell types identified in current data and the data of (Laurenti et al., 2015). f, Heatmap of enrichment score between cell types identified in current data and the data of (Ng et al., 2009). g, Heatmap of enrichment score between cell types identified in current data and the data of (Milyavsky et al., 2010). The color keys from shallow to deep indicate the range of −log_10_-transformed *P* value calculated by Hypergeometric distribution test. The horizontal axis of each heatmap represents the cell type of current data after dimensionality reduction and cell cluster, and the vertical axis represents cell type notes in bibliography.

**Figure S4. PCA visualization of identified cell types and cell assignment of each cell type along the developmental trajectory.**

a, 3-dimension PCA visualization of 12 identified cell types based on PCC. b, Cell assignment of each cell type along the developmental trajectory.

**Figure S5. Expression of cell type-specific regulons of lymphoid, myeloid and erythroid lineage.**

a, Expression of MPP-specific regulons (*RFX2*_91g and *HOXB6*_91g). b, Expression of LMPP- and MLP-specific regulons (ZNF780F_12g and *ZNF21_88g*). c, Expression of CMP-specific regulons (*E2F1*_540g and *E2F8_294g*). d, Expression of GMP-specific regulons (*RFX8*_extended_81g and *CEBPD*_52g). e, Expression of MEP-specific regulons (*GATA2*_579g and *GATA1*_618g). f, Expression of ProB-specific regulons (*TCF4*_13g and *RXRB*_59g).

**Figure S6. The developmental routine of HSPCs and gene expression patterns of three lineages (HSCs/MPPs, lymphoid, and myeloid/erythroid cells).**

a, Trajectory tree showing the developmental routine from HSCs to progenitor cells and downstream myeloid, erythroid and lymphoid lineages. Different color represents different cell type. b, Relative gene expression patterns of three states (Fig. 5b-c) along the pseudo time. The horizontal axis represents the pseudo time, and the vertical axis represents relative gene expression levels.

