## Supplementary figures and images for "Functional stemness-related genes revealed by single-cell profiling of naïve and stimulated human CD34^+^ cells from CB and mPB"

### Figure S1

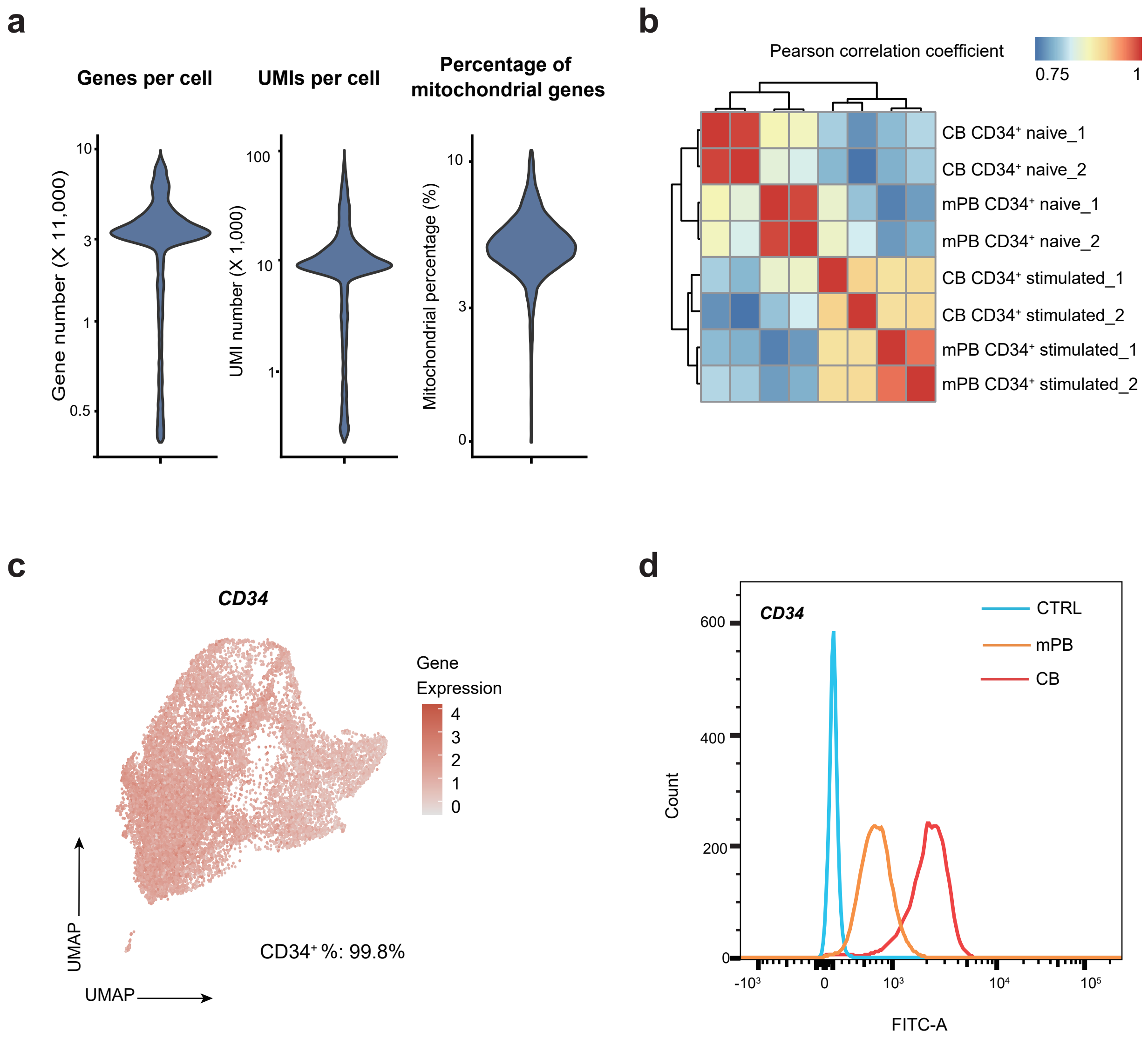

### Figure S2

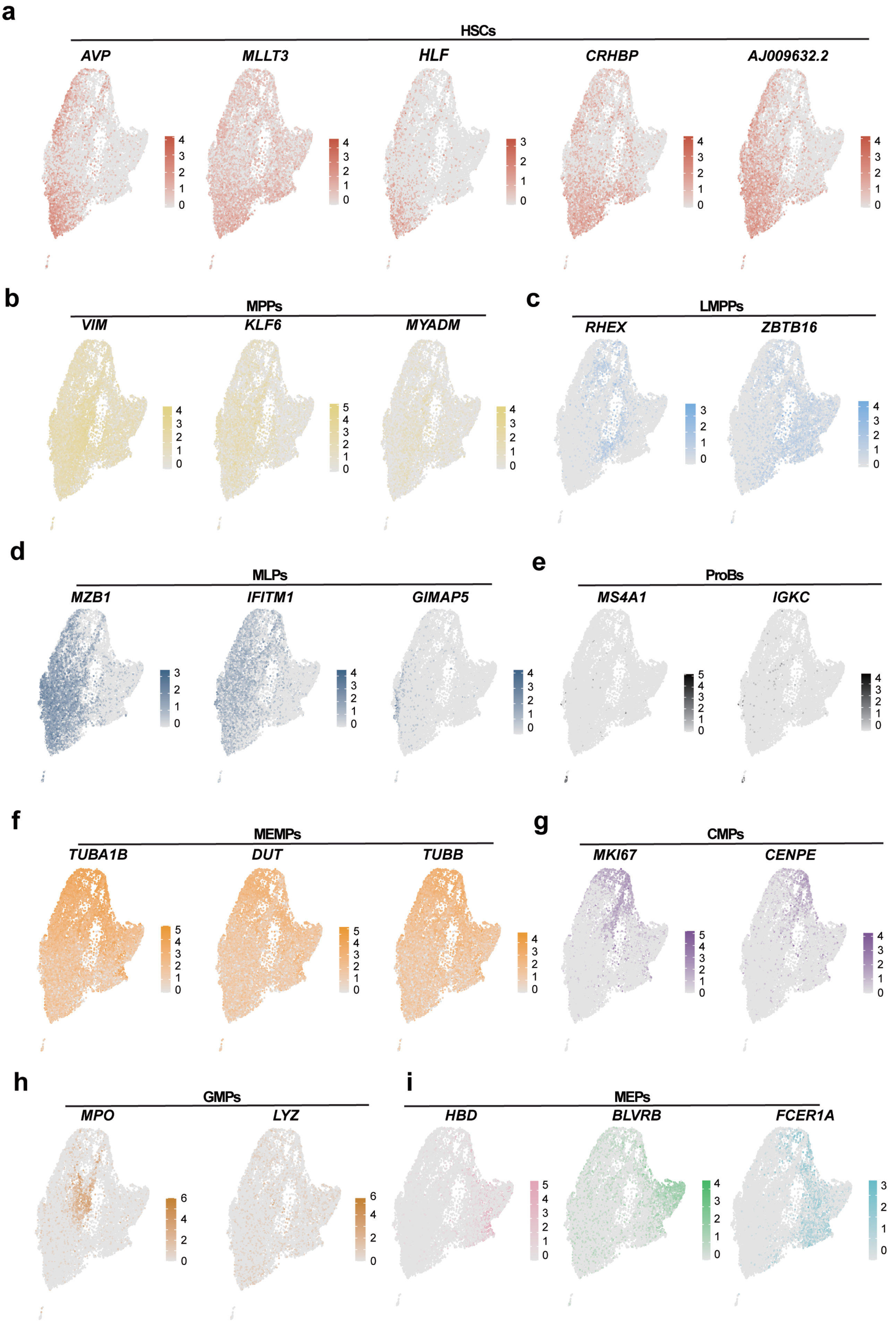

### Figure S3

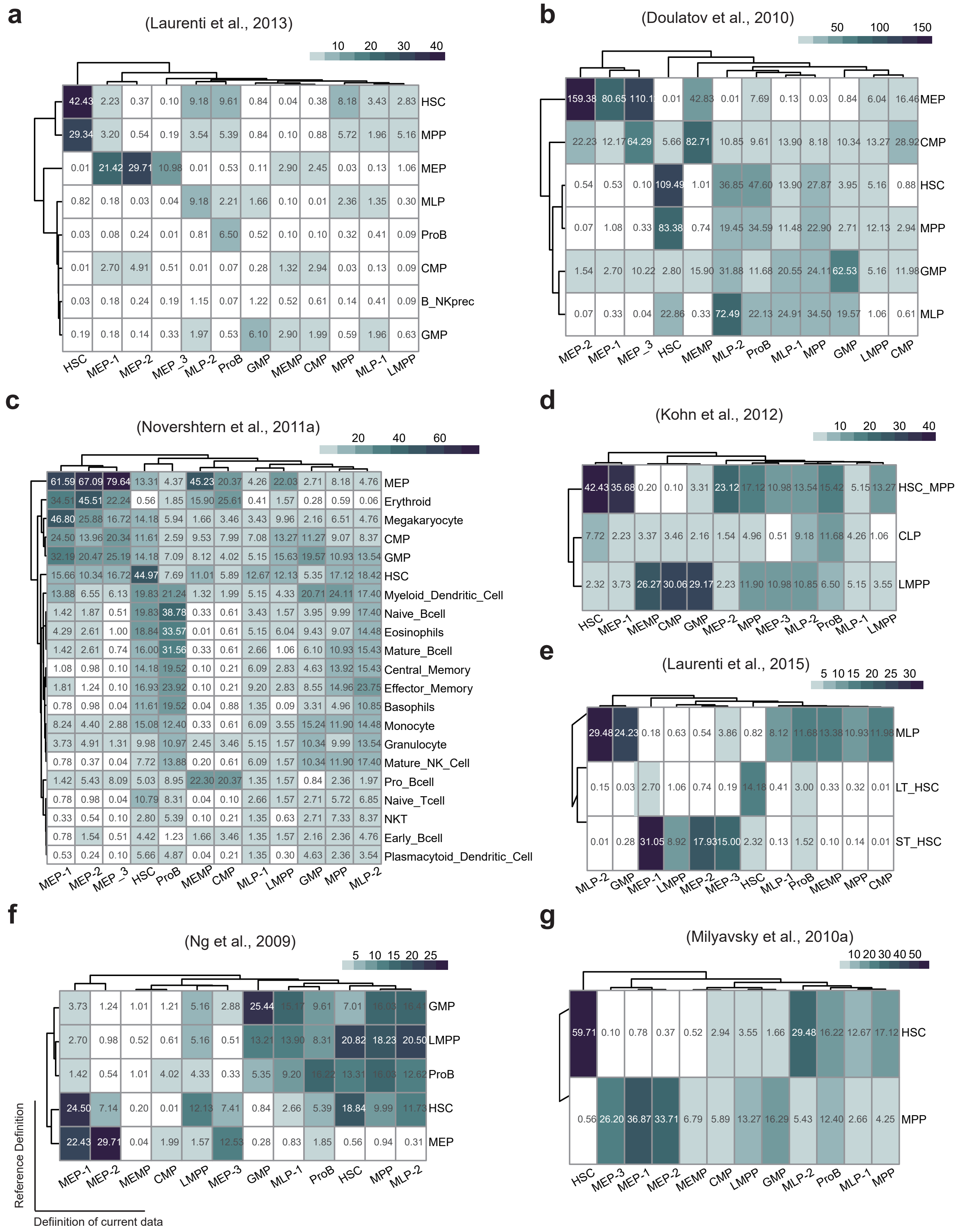

### Figure S4

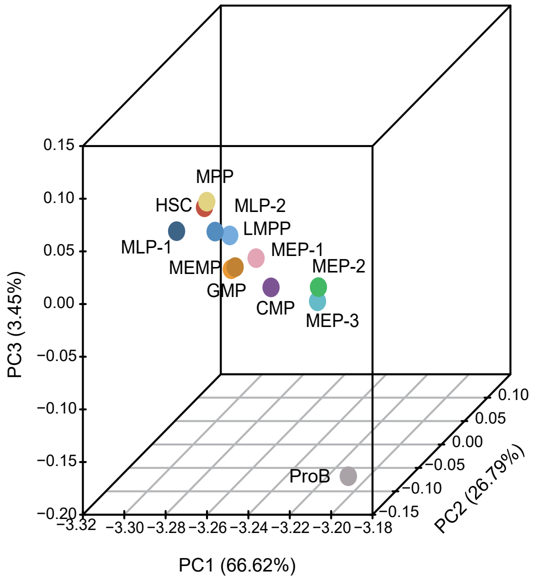

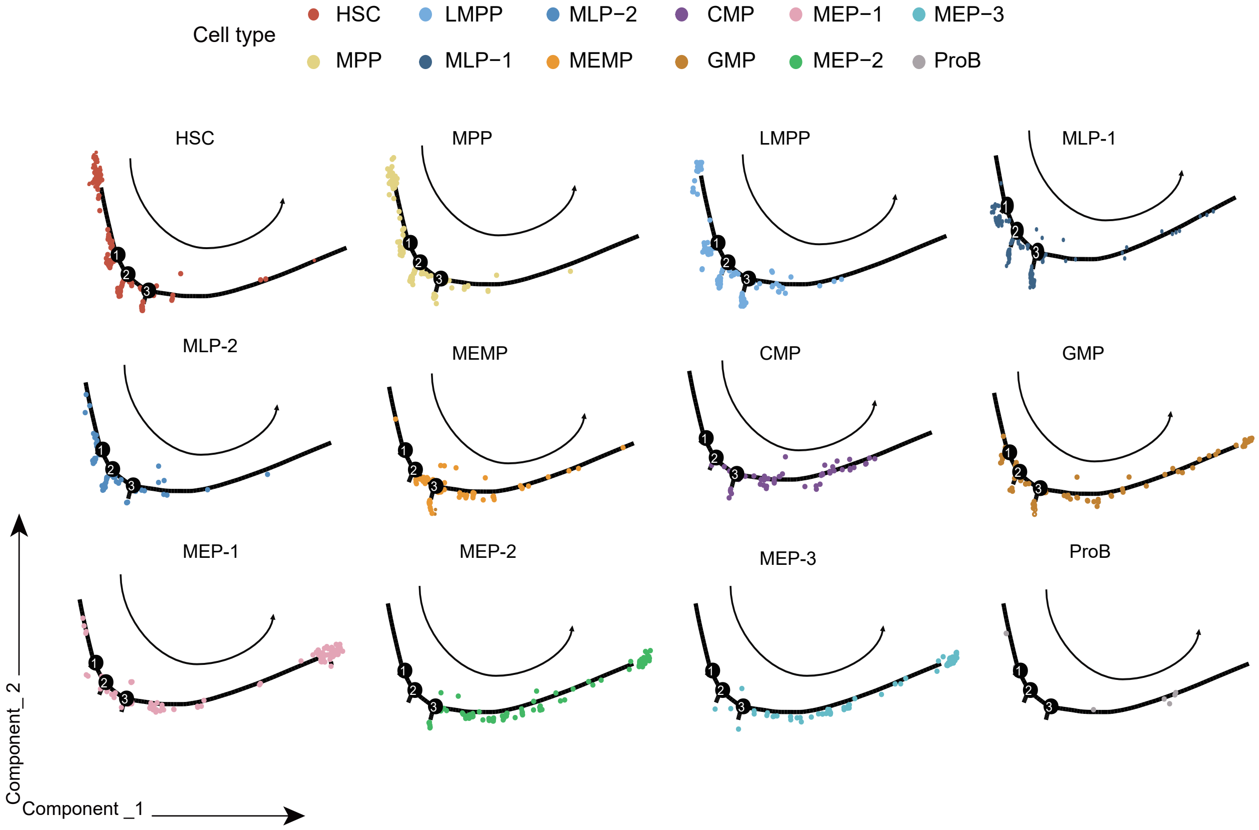

### Figure S5

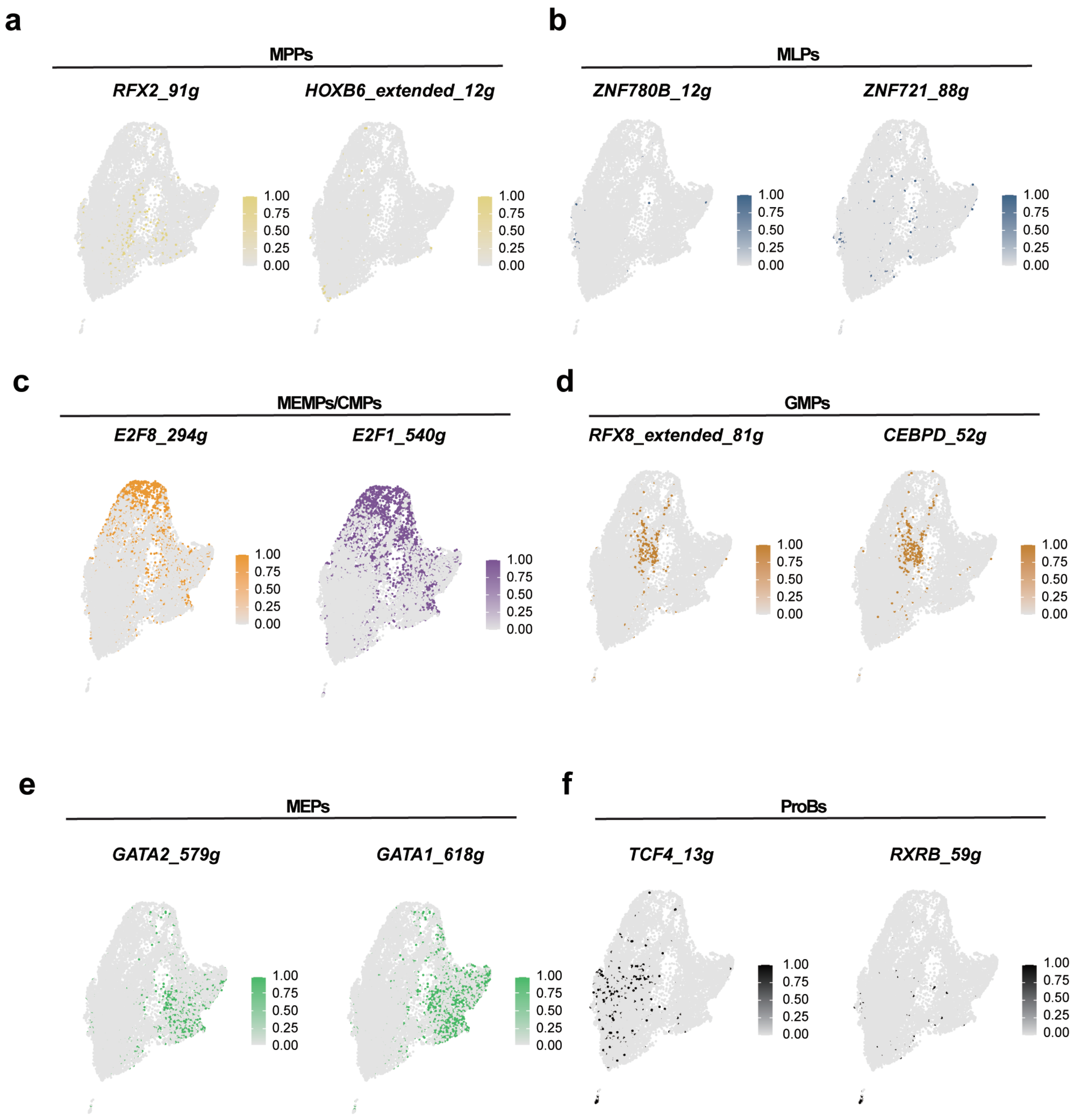

### Figure S6

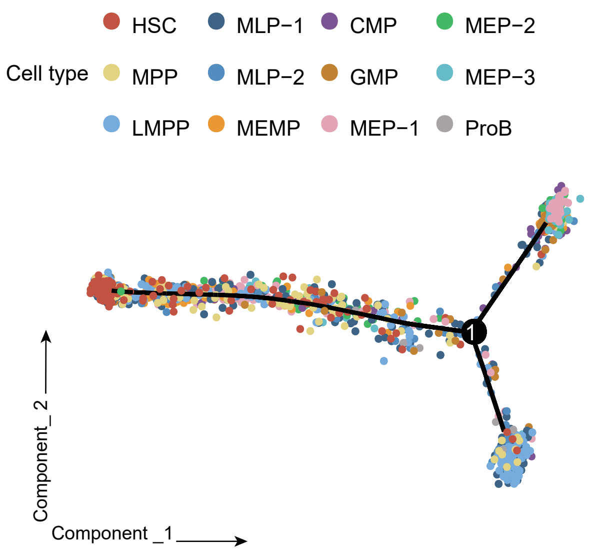

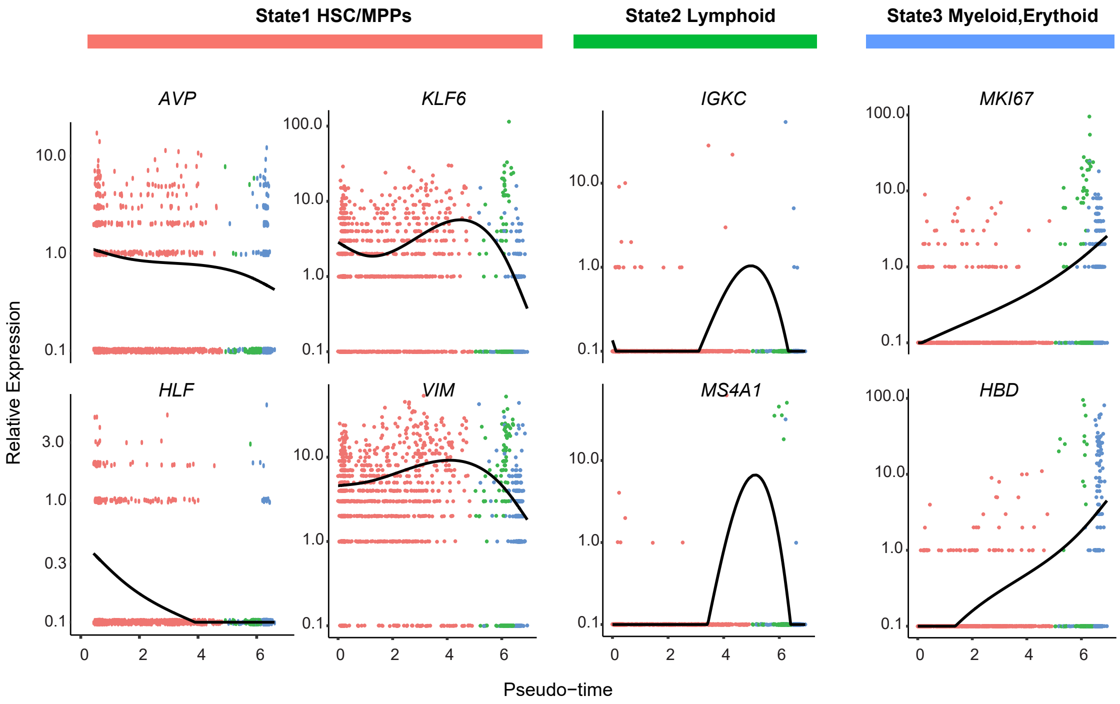
